# A transcription factor-pair work in concert to regulate gene expression across the life cycle of the pinewood nematode, *Bursaphelenchus xylophilus*

**DOI:** 10.64898/2026.06.24.734266

**Authors:** Madalena Mendonça, Anika Damm, Chongjing Xia, Cláudia S. L. Vicente, Sebastian Eves-van den Akker, Margarida Espada

## Abstract

The migratory endoparasitic pinewood nematode (PWN), *Bursaphelenchus xylophilus*, is the causal agent of pine wilt disease, causing significant economic and ecological losses in conifer forest ecosystems in Europe and Asia. Understanding the molecular mechanisms regulating PWN parasitism-related genes may lead to new sustainable solutions for control. Based on previous PWN transcriptomic datasets from the pre-parasitic and parasitic stages and from the pharyngeal gland cells (GC), an *in silico* analysis was performed to identify transcription factors (TF) highly expressed in the GC. Seven candidates TF genes were selected, and their spatial expression validated by *in situ* hybridisation. From those, two GC-expressed TFs, BXY_079 and BXY_022, each encoding zinc finger domains, were successfully knocked down by RNA interference. Transcriptomic data from silenced *BXY_079* and *BXY_022* TFs, analysed with existing life cycle specific transcriptomic data, showed that both TFs control genes expressed at similar times, by repressing male-related genes while activating genes expressed during the J3 and D3 stages, yet each represents the extreme of the others’ minor function. In addition to these common roles, BXY_079 also activates parasitism-related genes in the J2 stage. These BXY_079-activated parasitism-related genes predominantly encode proteins with lytic functions, including secreted peptidases and glycoside hydrolases. Consistent with their proposed role in parasitism, these genes are highly expressed during the parasitic juvenile stages and are likely involved in nematode feeding, tissue penetration, and migration within the host. In contrast, BXY_022 also represses the expression of several genes related to the reproduction system, such as major sperm proteins and cytosolic motility proteins, particularly in the adult male stage. Taken together, both dual-functional TFs work together, non-redundantly, to regulate gene expression across the life cycle, while each is additionally specialised to regulate diverse and distinct gene sets: ranging from genes implicated in lytic parasitic functions to sexual dimorphism.

## Introduction

Plant parasitic nematodes (PPN) are some of the most widespread and damaging pests in agronomy and forestry worldwide (Boyd *et al.,* 2013, Savary *et al*., 2019; Kantor *et al*., 2022). The pinewood nematode (PWN), *Bursaphelenchus xylophilus*, is a migratory plant endoparasite that has gained attention after its recognition as the causal agent of pine wilt disease, responsible for one of the most damaging threats to conifer forests (*Pinus* spp.) (Vicente *et al.,* 2012; Faria *et al.,* 2021; Kantor *et al*., 2022; Espada *et al.,* 2025). Listed as an A2 quarantine organism in Europe (EPPO, 2023), PWN represents a large threat of the forest ecosystems and wood industry, and the rapid spread of this disease has become a problem with the potential to cause significant economic and ecological damage (Mota and Vieira, 2008; Vicente *et al.,* 2012). PWN infects the host pine via transmission by the insect vector and, once inside, migrates into the resin canals, where it feeds and reproduces, consequently blocking the vascular system leading to wilting symptoms and tree death (Mota *et al*., 1999; Vicente *et al.,* 2012; Kantor *et al*., 2022). PWN has a life cycle which involves both mycophagous and phytophagous phases of development. This dual-feeding mode allows the PWN to survive and reproduce in the host at an advanced stage of the disease when the tree is declining and there is no healthy plant tissue available (Vicente *et al.,* 2012; Futai, 2013; Jones *et al.,* 2013; Vicente *et al*., 2025). PWN life cycle develops from the egg through four propagative juvenile stages (J1 to J4) to reach the adult stage. For transmission to a new host tree by the insect vector, PWN switches from propagative to the dispersal third-stage juvenile (D3), which subsequently moults into the fourth-stage dispersal juvenile (D4, dauer) (Wingfield *et al*., 1987; Hasegawa and Miwa, 2008; Futai, 2013; Tanaka *et al.,* 2019). Understanding the molecular mechanisms of PWN parasitism is essential to develop new control strategies due to the absence of efficient ones (Rutter *et al.,* 2014).

The interactions of PPN with the host are mediated by parasitism-related proteins (effectors), which are secreted into the host to promote disease (Haegeman *et al*., 2012; Bird *et al*., 2015; Gleason and Vieira, 2019). Most of these proteins are produced in the pharyngeal gland cells (GC), parasitism-specialised cells crucial in parasitism (Haegeman *et al.,* 2012; Gleason and Vieira, 2019). The identification and functional characterization of parasitism-related genes (effectors) is crucial for understanding nematode parasitism of plants, as functionally altering or blocking these genes can reduce the nematode’s ability to infect the host (Maier *et al*., 2013; Gleason and Vieira, 2019). Espada *et al*. (2018) were able to identify a non-coding DNA motif-STATAWAARS that is associated with the promoter region of a large set of PWN-secreted parasitism-related genes. Using this strategy, Espada *et al*. (2018) were able to use this motif as a global genome predictor of novel parasitism-related genes, providing a new set of candidates specific to the PWN, which can be useful to explore as targets for nematode resistance. Additionally, similar DNA motifs were previously discovered and associated with effector gene GC-expression in non-related PPNs (Eves-van den Akker *et al.,* 2016, Vieira *et al.,* 2021, Da Rocha *et al.,* 2021, Pellegrin and Damm *et al.,* 2025). The discovery of these different promoter elements in a large set of both migratory and sedentary nematodes, suggest that secretion might be orchestrated by regulatory protein(s) specific to the GC that can bind this sequence to control the expression of genes, such as transcription factors (TFs) (Eves-van den Akker *et al.,* 2016, Espada *et al*., 2018). Identifying the TFs involved, as well as the upstream signals that activate them, is therefore fundamental to understand the effector gene regulation. As an example, a subventral gland cells regulator transcription factor has been found in the cyst nematode *Heterodera schachtii,* that directly binds to parasitism gene promoters and activates their transcription during the early stages of parasitism. The knockdown of the regulator reduced the expression of parasitism-related genes and had a major impact on the ability to infect the host (Pellegrin and Damm *et al.,* 2025).

Here we identified and functionally characterised two zinc finger TFs, BXY_079 and BXY_022, which are spatially expressed in the GC, for the first time in PWN. We investigated their role by RNA interference and compared the silenced transcriptomic data with existing life cycle expression data (Tanaka *et al.,* 2019). Our results indicate that both TFs may work together, non-redundantly, to regulate gene expression by activating and repressing across the PWN life cycle, while each is additionally specialised to regulate diverse and distinct gene sets: ranging from genes implicated in lytic parasitic functions to sexual dimorphism.

## Results and discussion

### Identification of two gland-cell expressed transcriptions factors, BXY_079 and BXY_022

Based on previous PWN transcriptomic datasets from the pre-parasitic and parasitic stages (Espada *et al.,* 2016), as well as transcriptomics of isolated GC (Espada *et al*., 2018), a comparative *in silico* analyses was performed to identify GC enriched transcripts. Seven genes encoding putative transcription factors were highly represented in the gland cell libraries than the whole worm libraries, by a factor of 2- to 5-fold (data not shown), suggesting that they may be resident in, and associated with the regulation of, gland-cells. From the seven candidate TF genes identified, four were subjected to RNAi-mediated silencing. Two of these candidates (BXY_079 and BXY_022) were amenable to both *in situ* hybridisation, which supported enriched expression profiles in the gland cells compared to the whole body (Figure 1A, B) and RNA interference (which enables interrogation of molecular function, Figure 1A, B). Both TFs encode zinc finger domain-containing proteins but from different families: BXY_079 encodes a protein most similar to the C_2_H_2_ family which is commonly associated with DNA-binding (Brayer *et al*., 2008), while BXY_022 encodes a protein most similar to the RING type C_3_HC_4_ family which participate in a wide range of cellular processes, such as transcriptional regulation, signal transduction, and recombination (Wu *et al*., 2014). The RING domain may also function as a protein-protein interaction module and is commonly associated with ubiquitin ligase activity, mediating ubiquitination-dependent regulatory pathways (Stone *et al*., 2005; Wu *et al*., 2014). In eukaryotes, these families of zinc finger domain-containing proteins perform a wide range of biological functions, including DNA and RNA binding, transcriptional regulation, apoptosis control, protein folding, and lipid interactions (Laity *et al*., 2001; Brayer *et al*., 2008). To interrogate their molecular function, we used RNAi-coupled RNA-seq to measure whether and how BXY_079 and BXY_022 contribute to transcriptional regulation in PWN.

**Figure 1.**
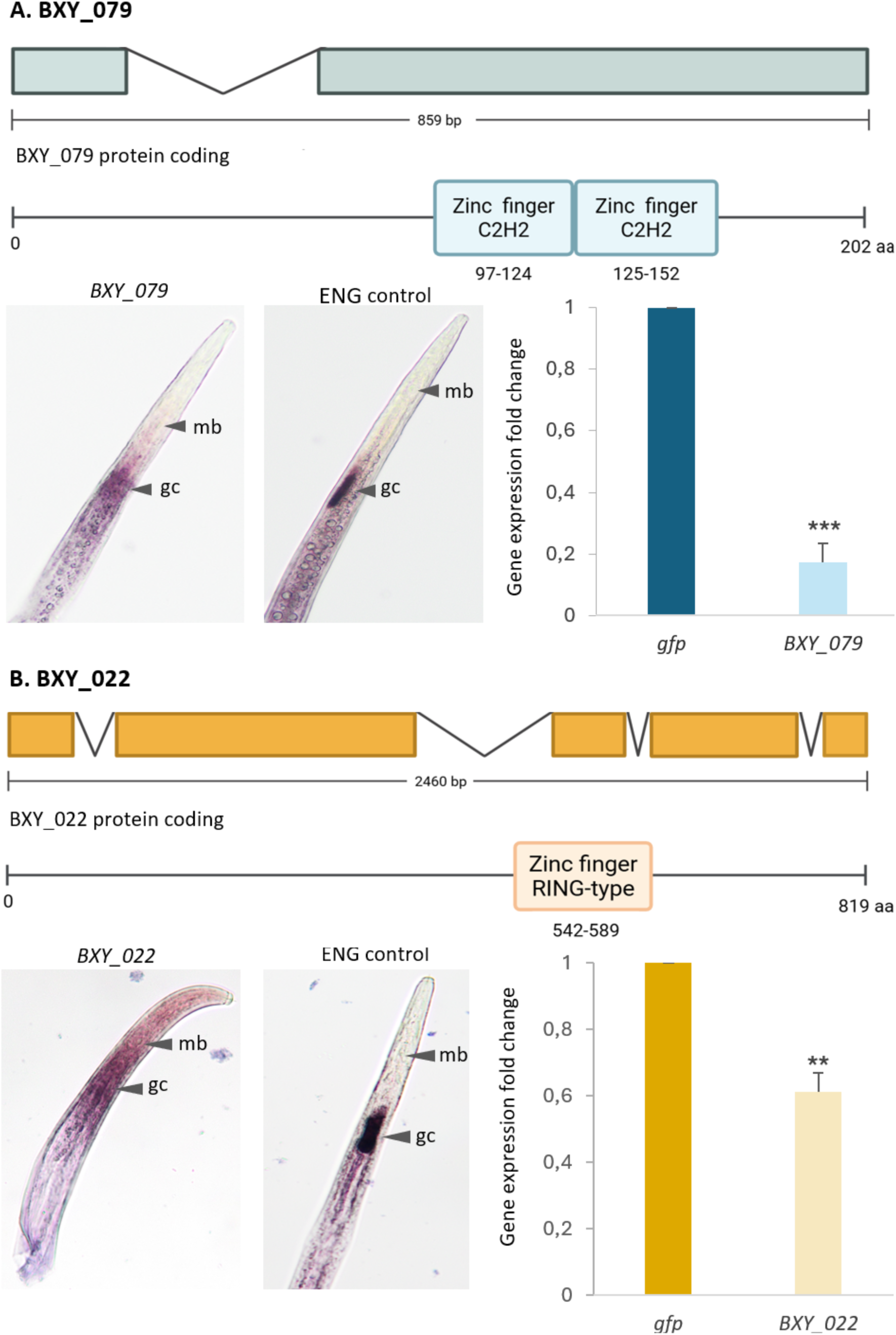
Characterisation of zinc finger transcription factors. A) BXY_079 and B) BXY_022. Representation of BXY_079 and BXY_022 domains, intron (lines) and exons (coloured boxes) (Biorender.com). Gene expression following RNAi knockdown compared to a *gfp* control, determined by RT-qPCR and normalised using 2^-ΔΔCt^ method (Livak & Schmittgen, 2001). Asterisks indicate significantly different treatments compared to the respective *gfp* control at (*) *p* < 0.05, (**) *p* < 0.01, and (***) *p* < 0.001 (Tukey’s test), which were significantly different compared with the control treatment. *In situ* hybridisation of BXY_079 and BXY_022 TFs candidate genes show the (purple) signal in the GCs area (ENG was used as positive control). The different nematode sections of the body are labelled as follows, gc: pharyngeal gland cells; mb: median bulb.

### BXY_079 activates the expression of candidate parasitism-related genes

In *BXY_079*-silenced nematodes, transcriptomic analysis identified 563 differentially expressed genes (DEG) (|log2FC| ≥ 0.5, Padj ≤ 0.05) compared to the *gfp* control. Given that under *BXY_079*-silenced conditions, the dominant majority of DEGs are downregulated (83%, Figure 2A), *BXY_079* is predominantly a positive regulator of gene expression under native conditions. Interestingly, BXY_079-activated genes are enriched in putatively secreted proteins (115/467, 24.6% vs ∼12% (2104/17560) for genome PRJEA64437 as a whole (p=1.37546E-08 hypergeometric test; Figure 2A). Of these, nearly a third are associated with lytic functions, including cysteine, aspartic, and serine peptidases, as well as glycoside hydrolases (GH). This observation is consistent with gene ontology (GO) enrichment analyses, which revealed significant enrichment of proteolytic functions, including aspartic-type peptidase activity, aspartic-type endopeptidase activity, and endopeptidase activity, as well as proteins associated with lysosomal/lytic vacuole compartments, suggesting the presence of degradative enzymes involved in proteolysis and intracellular digestion (Supplementary Figure S1). Many BXY_079-activated transcripts, including those which are predicted to encode such lytic functions, are themselves more highly expressed in previous gland cell sequencing libraries than whole body libraries (Figure 2B), as is *BXY_079* itself.

**Figure 2.**
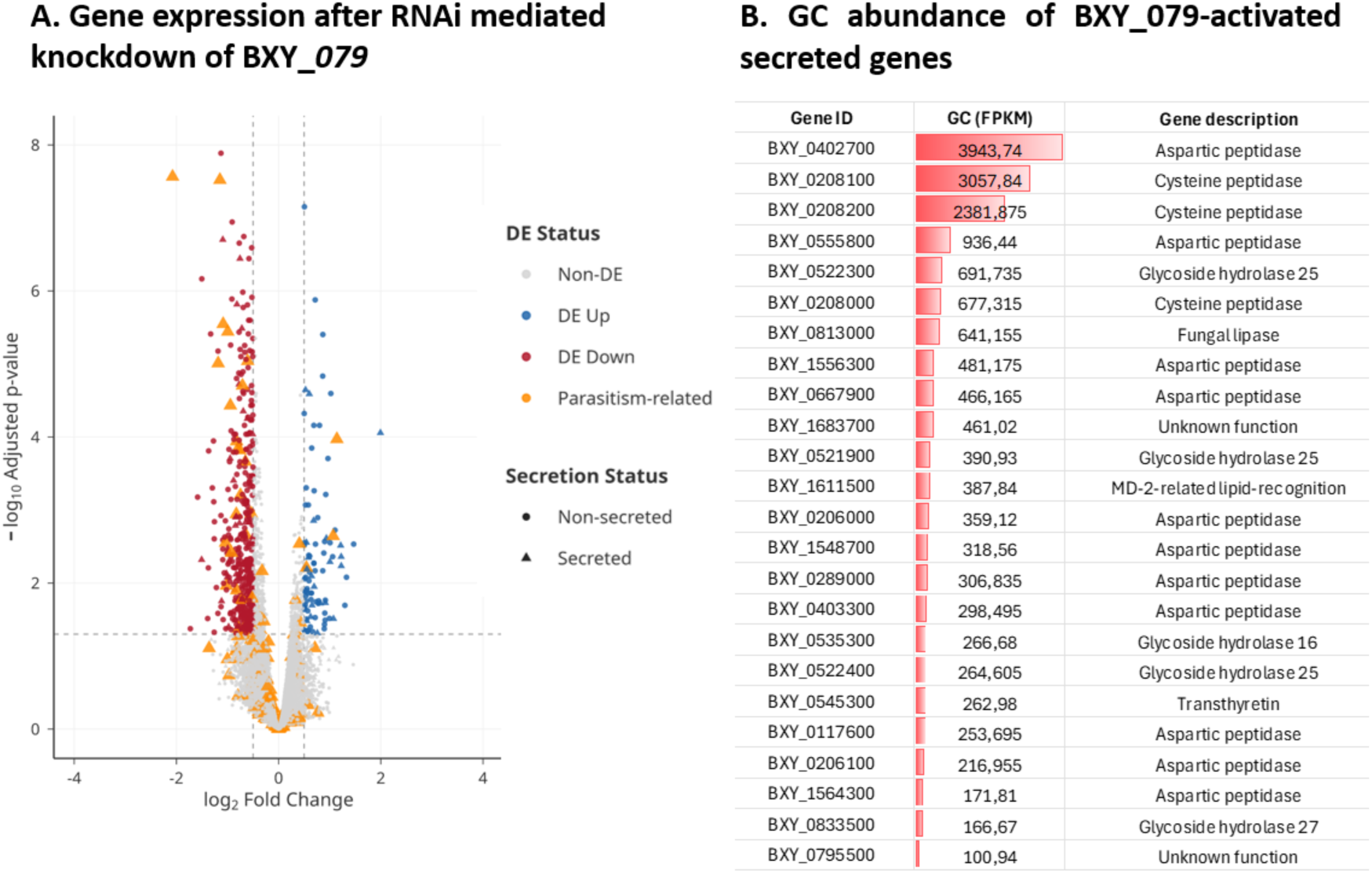
Differential gene expression after *BXY_079* knockdown and GC expression. A) Volcano plot of PWN gene expression following *BXY_079* knockdown vs. *gfp* control. B) Bars representing the abundance expression profile of the several activated secreted DEG above 100 using the fragment per kilobase of transcript per million mapped reads (FPKM) values of the GC library from Espada *et al*. 2018.

We therefore also validated *BXY_079-*regulation, and gland cell expression, of nine BXY_079-activated transcripts, including those associated with lytic functions and others (Figure 3A and 3B).

**Figure 3.**
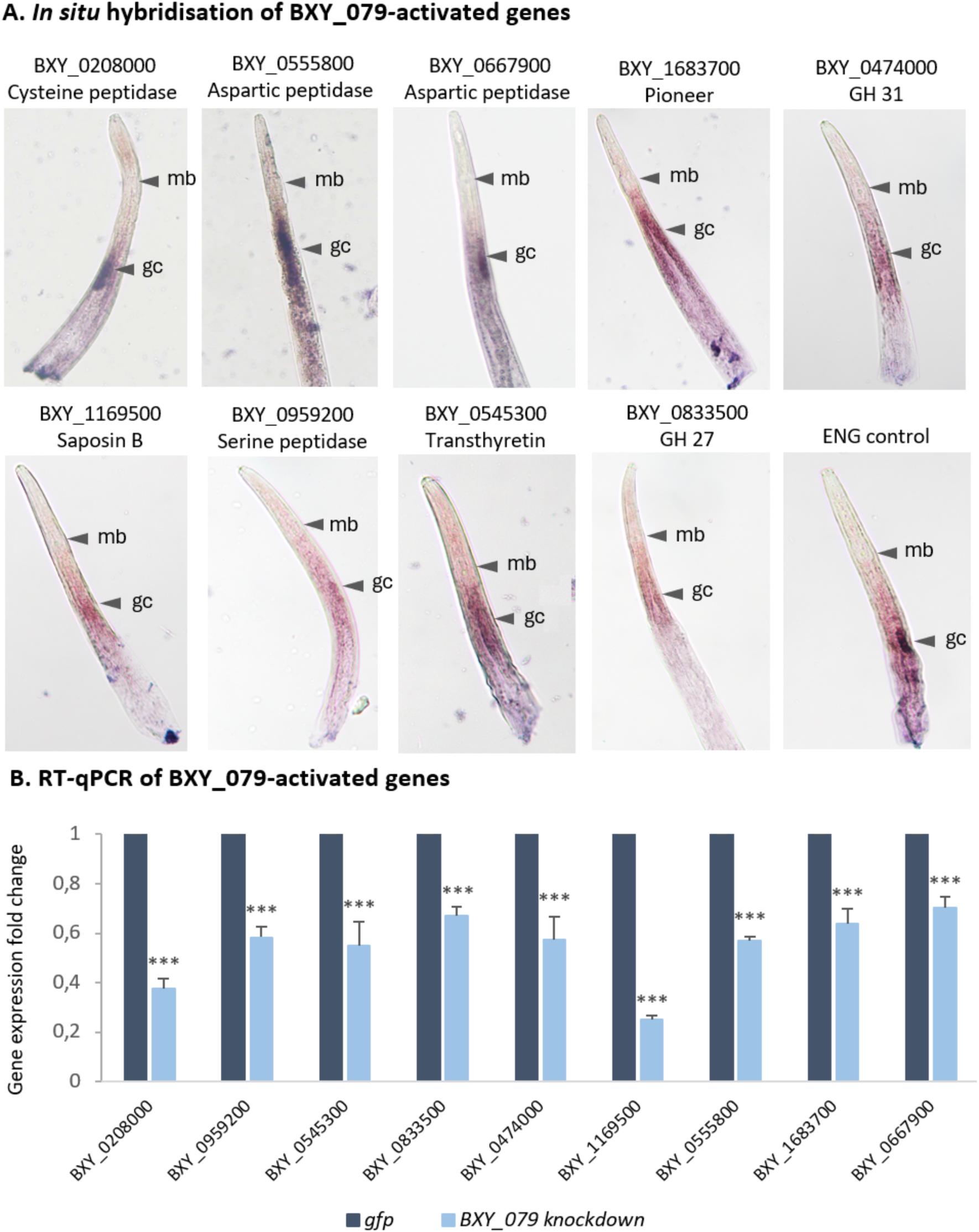
Spatial expression and relative gene expression of BXY_079-activated genes. A) *In situ* hybridisation of nine BXY_079 regulated parasitism-related DEGs candidates. B) Relative gene expression of BXY_079-activated genes following *BXY_079* knockdown vs. *gfp* control was determined by RT-qPCR and normalised using 2^-ΔΔCt^ method (Livak & Schmittgen, 2001). Asterisks indicate significantly different treatments at (***) *p* < 0.001 (Tukey’s test), which were significantly different compared with the *gfp* control treatment (zero).

Cross-referencing BXY_079 activated genes with existing life cycle expression data (Tanaka *et al.,* 2019) indicates that BXY_079-activated lytic functions are predominantly expressed during juvenile stages (Figure 4A), which may reflect their involvement in early host invasion and tissue degradation necessary for successful parasitism, and is consistent with the expression pattern of *BXY_079* itself: J2, J3, and J4 (Figure 4B). Interestingly, while BXY_079 is dominantly a positive regulator of gene expression, it also represses some genes. BXY_079-repressed genes are not significantly enriched in any GO terms, but they are enriched in genes expressed at the male life stage, and the D4 (Figure 4C).

**Figure 4.**
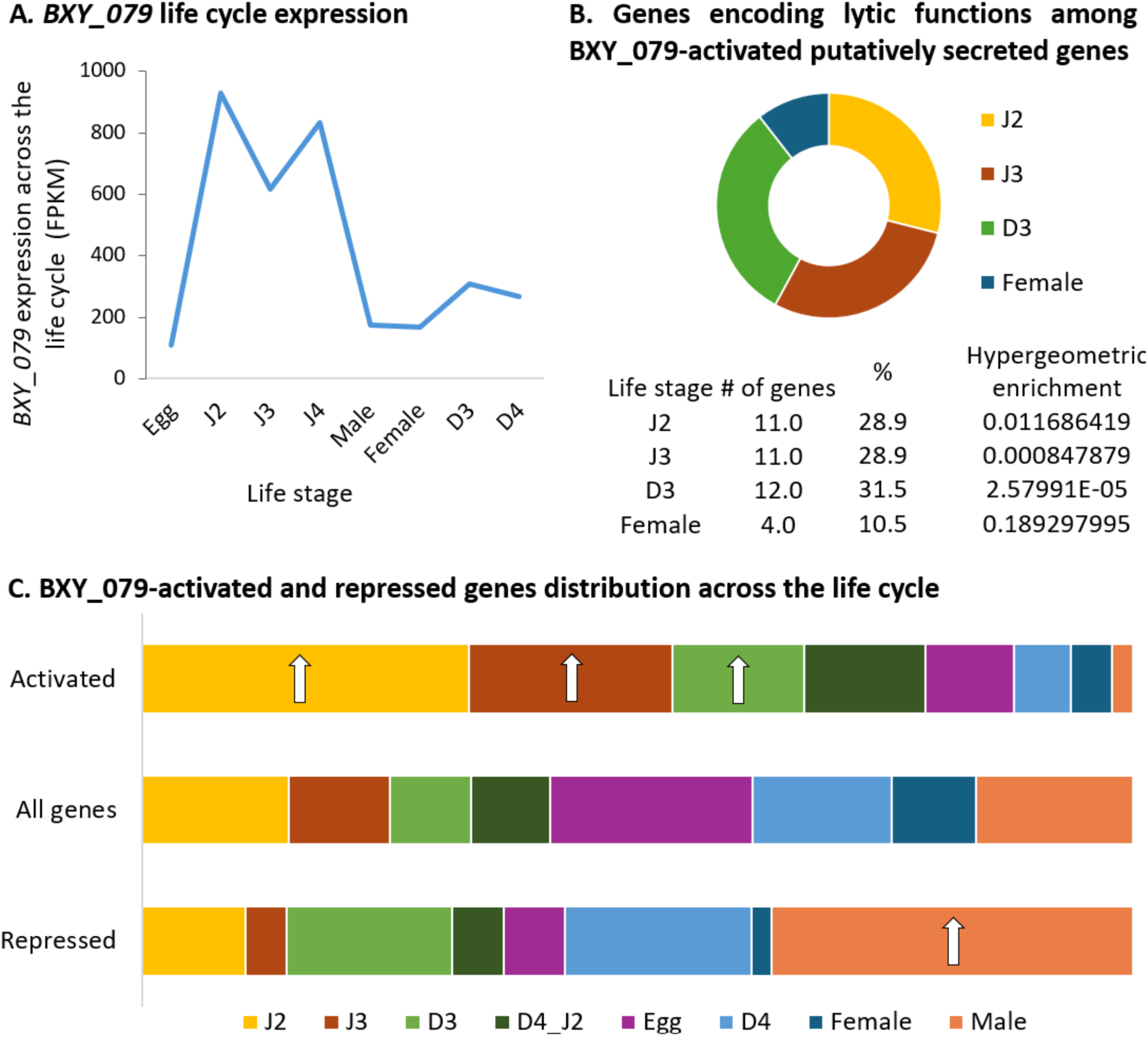
BXY_079. A) *BXY_079* gene expression across the nematode life cycle. B) Pie chart showing the distribution (%) of genes encoding lytic functions among BXY_079 activated genes across different life stages of the PWN. C) BXY_079-activated and repressed genes across the PWN life cycle existing data. Arrows indicate the life stages showing significant enrichment.

Peptidases are essential enzymes that hydrolyse peptide bonds and play key roles in fundamental physiological processes, such as embryogenesis and cuticle remodelling during larval development, as well as in parasitic strategies, including host tissue invasion, nutrient acquisition through protein degradation, and evasion of host immune defenses (Kikuchi *et al*., 2011; Espada *et al*., 2025). Several GH families were also identified, including GH16 (xyloglucan:xyloglucosyltransferases), GH31 (α-Glucosidases), and GH27 (α-Galactosidases), with the GH25 (lysozyme) family, being the most abundant (Rai *et al*., 2015). These GH enzymes are involved in carbohydrate metabolism and constitute a major group of cell wall-degrading enzymes that hydrolyse polysaccharides of plant and fungal cell walls (den Brink & de Vries, 2011; Cardoso *et al.,* 2016). Lysozyme protein was found to be highly upregulated during infection and spatially expressed in the GC, suggesting a potential role in parasitism in PWN (Espada *et al*., 2016). Moreover, these enzymes may contribute to the digestion of host proteins (Espada *et al*., 2016) and could also have an antibacterial activity by of cleaving the peptidoglycan component of bacterial cell walls, as has been demonstrated in *Caenorhabditis elegans* (Nicholas & Hodgkin, 2004). BXY_0555800 (aspartic peptidase) and BXY_1236200 (histidine acid phosphatase), downregulated DEG, were found be increased proteins in PWN secretome under the *P. pinaster* stimulus (Silva *et al*., 2021). BXY_079 positively regulates effector genes predicted in the literature including a GH16 (Kikuchi *et al.,* 2005), GH25, glutathione S-transferase and a cysteine peptidase protein (Espada *et al*., 2016).

### BXY_022 represses the expression of reproduction system genes

In BXY_*022*-silenced nematodes, transcriptomic analysis identified 479 DEGs (|log2FC| ≥ 0.5, and *P*adj ≤ 0.05) compared to the *gfp* control (Fig.5A). Given that under *BXY_022*-silenced conditions, the dominant majority of DEGs are upregulated (69.10%), BXY_022 is - contrary to BXY_079 *- a* predominantly negative regulator of gene expression under native conditions. BXY_022-repressed genes are not associated with putatively secreted proteins (12.68%; enrichment of 4.31%), and BXY_022 is more stably expressed across the life cycle, with marked but only minor increase at the D3 dispersal stage followed by a decrease at D4 (Fig.5B). However, BXY_022-repressed genes are significantly enriched in GO terms (Supplementary Figure S2) associated with protein phosphorylation regulation, particularly serine/threonine phosphatase activity, and transmembrane transport of amino acids and organic compounds. RT-qPCR validation was performed to confirm the negative relationship between BXY_022 and genes encoding these functions, consistent with the RNA-seq results (Fig.5C). Among the 289 DEGs encoding non-secreted proteins, several genes are associated with reproduction and development such as major sperm proteins (MSPs), cytosolic motility proteins (CMP), sperm meiosis PDZ domain-containing proteins (SMZ) and the only two interactor of ZYG-11 genes in the whole PWN genome. MSPs play a central role in the motility machinery that enables the crawling movement of nematode sperm (Roberts *et al*., 2005). BxMSP10, identified as repressed DEG, has previously been shown to be specifically expressed in the seminal vesicle of adult male PWN and essential for reproduction and egg hatching (Huang *et al*., 2019). MSPs, in complex with cytosolic motility proteins, allow motility in nematodes (Grant *et al.,* 2005) and sperm meiosis PDZ domain-containing proteins have recently been identified in *Caenorhabditis elegans* as key regulators of nematode spermatogenesis, promoting MSP filament assembly and the formation of fibrous body-membranous organelles, structures essential for sperm development (Peng *et al*., 2025). Moreover, interactor of ZYG-11 proteins has been identified in *C. elegans*, as a substrate-recognition subunit for a CUL-2 based complex that regulates cell division and embryonic development (Vasudevan *et al*., 2007), meaning that these genes altogether are likely contribute to essential processes in PWN reproduction and development. Consistent with this, many BXY_022-repressed genes are themselves expressed in the male stage (p-value=1.7813E-157, 81.3%) (Figure 5D). Taken together, the coordinated expression of MSPs, CMPs, SMZs, and ZYG-11 interactors may indicate that these non-secreted proteins are critical for male fertility and embryonic development in PWN, highlighting their potential stage- and sex-specific functions in reproduction. These results suggest that BXY_022 probably acts by repressing the expression of genes related with reproduction and development in PWN.

**Figure 5.**
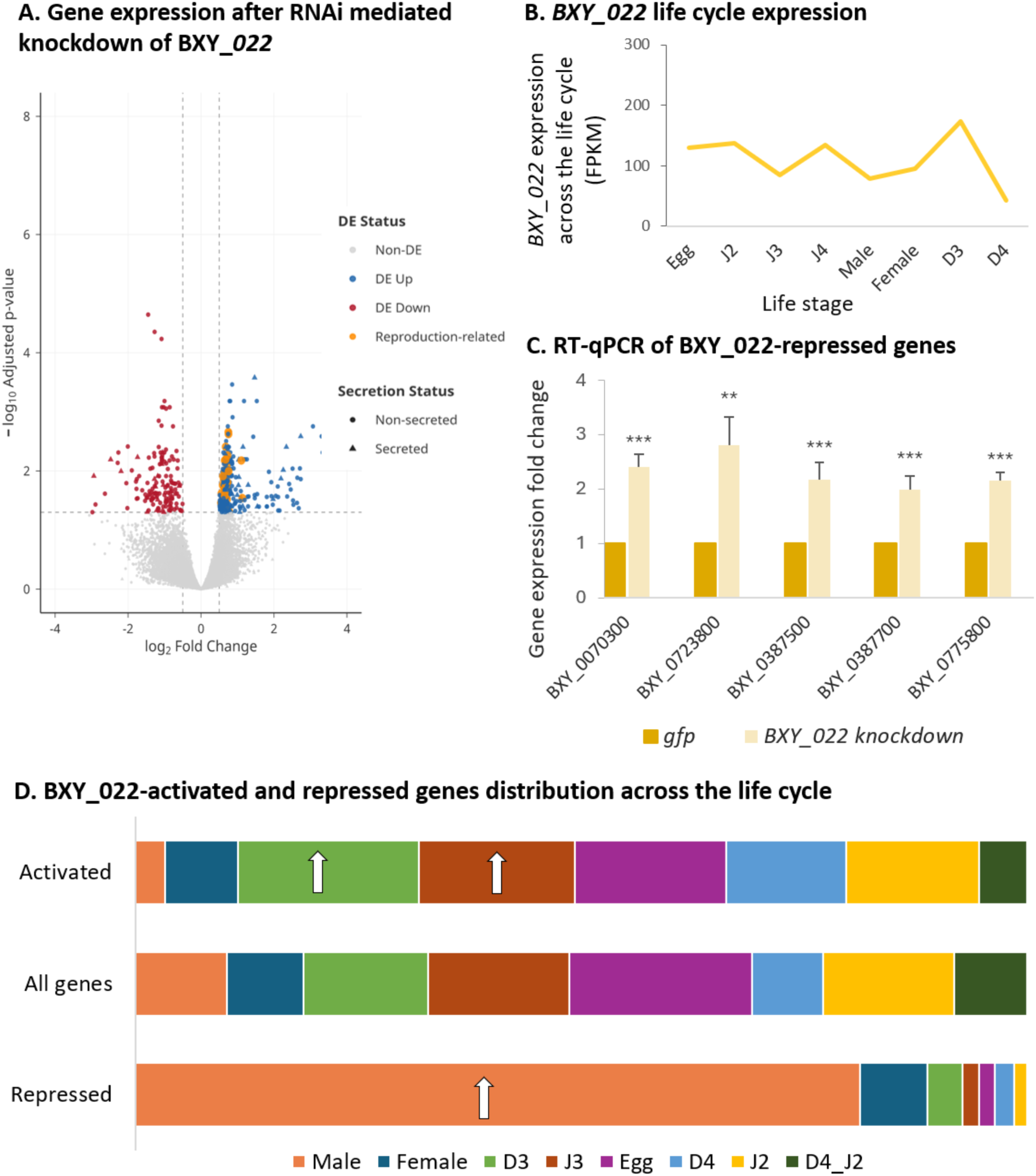
BXY_022. A) Volcano plot of PWN gene expression following *BXY_022* knockdown vs. *gfp* control. B) *BXY_022* gene expression across the nematode life cycle (data from Tanaka *et al*., 2019). C) Relative gene expression of BXY_022-repressed genes following *BXY_022* knockdown vs. *gfp* control was determined by RT-qPCR and normalised using 2^-ΔΔCt^ method (Livak & Schmittgen, 2001). Asterisks indicate significantly different treatments at (**) *p* < 0.01, and (***) *p* < 0.001 (Tukey’s test), which were significantly different compared with the *gfp* control treatment (zero). D) BXY_022-activated and repressed genes across the PWN life cycle existing data. Arrows indicate the life stages showing significant enrichment.

Interestingly, and conceptually similar to BXY_079, while BXY_022 is dominantly a negative regulator of gene expression, it also activates some genes. These BXY_022-activated genes are also not significantly enriched in any GO terms but do disproportionately contain genes expressed in J3 and D3 stages under native conditions (p-value=0.002233171, 17.6% and 1.88017E-06, 20.3%, respectively).

### BXY_079 and BXY_022 repress genes in adult males and activate genes in juvenile stages of the PWN life cycle

The comparative analysis between the transcriptomic data from silenced TFs with existing PWN life cycle specific transcriptomic data (Tanaka *et al*., 2019) reveal a broadly similar pattern for both BXY_079 and BXY_022, where each represents the extreme of the others’ minor function. As evidence, BXY_079 and BXY_022-activated genes contain, respectively, 2.0 and 1.6 times as many genes expressed at the J3 stage as compared to a random distribution (p-values= 1.00042E-11 and 4.48342E-05 respectively), as well as 1.7 and 2.5 times as many genes expressed at the D3 compared to random (p-values=0.002233171 and 1.88017E-06 respectively). On the other hand, BXY_079 and BXY_022-repressed genes contain 5.1 and 2.3 times as many genes expressed at the male stage as compared to random (p-values= 1.7813E-157 and 4.45021E-07 respectively). Taken together with the predominantly activating function of BXY_079 and the predominantly repressing function of BXY_022, it can be concluded that both BXY_079 and BXY_022 activate genes in the J3 and D3 stage, and repress genes in the male stage: but BXY_079 is numerically dominant in the former while BXY_022 is numerically dominant in the later (Figure 6A).

**Figure 6.**
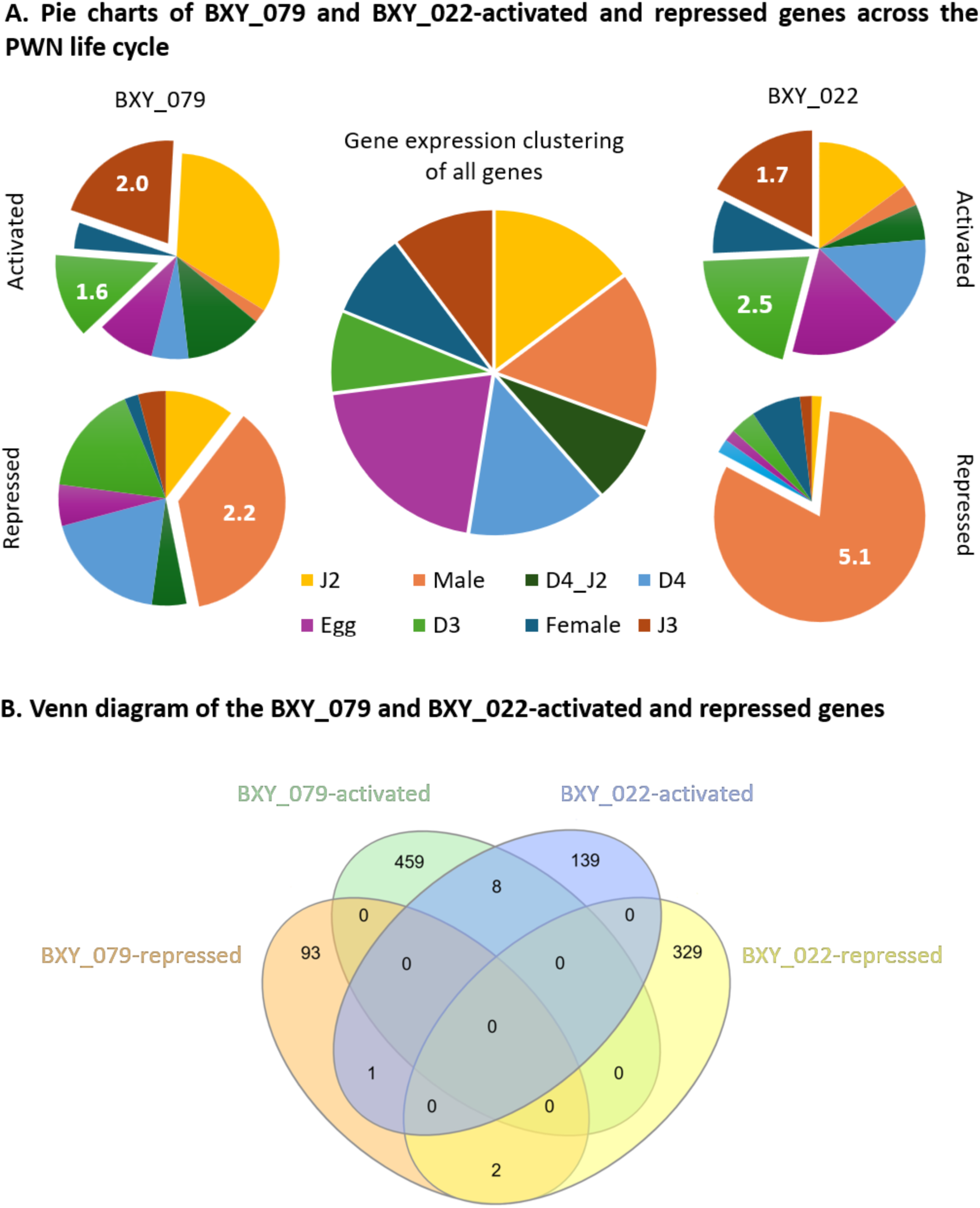
Gene expression distribution across the life cycle of PWN among activated and repressed genes regulated by BXY_022 and BXY_079 A) Pie charts illustrate the proportion of activated and repressed genes whose expression is associated with life cycle stages, with fold change in the most relevant stages. B) Venn diagram showing the overlap between activated and repressed genes regulated by the transcription factors BXY_022 and BXY_079 (Herberle *et al*., 2015).

Interestingly, of the approximately 1000 genes regulated by both genes combined, only eleven are co-regulated (Figure 6B). Eight are commonly activated, two are commonly repressed, and one is discordantly activated (Figure 6B). This indicates that although BXY_079 and BXY_022 are broadly consistent in their control of genes expressed at the same time, they regulate largely non-overlapping gene sets.

It is well established that TFs are generally classified as activators or repressors based on their effect on target gene expression, although some TFs are dual-functional and can act as either activators or repressors depending on the molecular context, or downstream/indirect targets (Boyle and Després, 2010; Martinez-Corral *et al*., 2025). Our results indicate that both TFs function as dual regulators, capable of both activating and repressing genes, but BXY_079 predominantly functions as an activator and BXY_022 mainly acts as a repressor. Moreover, genes are typically transcriptionally activated in stages where their function is required and repressed when it is not, resulting in increased transcript abundance in the relevant stages (Kikuchi *et al.,* 2017). These results support the existence of a TF duet and suggest that BXY_079 and BXY_022 act in concert, likely with other partners, at the same life stages, while regulating largely distinct gene sets. Rather than functioning redundantly, they may work together in parallel, each with its specialised function to contribute to the overall stage specific expression of the PWN life cycle. This coordinated yet non-redundant activity highlights their combined role in the transcriptional control of key biological processes in the PWN. These findings identify BXY_079 and BXY_022 as key regulators of stage-specific gene expression in PWN and highlight them as promising targets for novel biocontrol strategies. Their roles in developmental and parasitism-related pathways may make them attractive candidates for both RNAi-based approaches and the development of small-molecule inhibitors aimed at disrupting essential regulatory networks required for nematode survival and pathogenicity.

## Materials and Methods

### Nematode culture conditions/biological material

The isolate of PWN used in this study, reference culture BX013.003 (N 39◦43’338”, W 9◦01’557”) from INIAV Nematology collection (INIAV I.P., Oeiras, Portugal), was cultured at Nemalab, University of Évora, Portugal. The nematodes were maintained in Erlenmeyer flasks containing *Botrytis cinerea* on barley seeds at 25°C (Evans, 1970) for approximately 10 days. Mixed life-stage nematodes were extracted using the modified Baermann funnel technique (Southey, 1986), followed by sieving (38 μm) and collected in distilled water.

### Identification of the candidate transcription factors by *in silico* analysis

Transcriptomic datasets previously generated from two different libraries, whole PWN body (pre and parasitic stages at different time points) and GC (Espada *et al.,* 2016, 2018), were analysed to identify a large set of genes with potential relevance for parasitism of the PWN and potential regulatory elements(s) GC-expressed. We conducted a comparative *in silico* analyses on the GC versus whole nematode transcriptomic datasets and prioritised seven candidate genes that: are putatively encode for TFs; and had 2 to 5 times-fold enriched profiles in the GC compared to the whole nematode body transcripts. From the seven candidate TF genes identified, four were subjected to RNAi-mediated silencing. Gene knockdown efficiency was successful for two target genes, BXY_079 and BXY_022, that were further characterised. The spatial validation of the TFs was performed as described in the *in situ* hybridisation section.

### RNA interference

The double stranded RNA (dsRNA) for *BXY_079* was synthesised using the MEGAscript RNAi Kit (Thermo Fisher Scientific, USA) according to the manufacturer’s instructions. The small interference RNA (siRNA) for *BXY_022* was synthesised using the Silencer siRNA Construction Kit (Thermo Fisher Scientific, USA) according to the manufacturer’s instructions. The quantity and quality of the dsRNA were assessed by an ND-2000/2000c NanoDrop spectrophotometer (Thermo Fisher Scientific, USA) and gel electrophoresis. Mixed life stages-PWN were soaked in the silencing mix with 50mM octopamine and 3mM spermidine using 1µg/μL *BXY_079* dsRNA and *BXY_022* siRNA and *gfp* used as a negative control, for 24 hours, 70 rpm, at 25°C in the dark. The primers used for this analysis are shown in Supplementary Table S1. Silenced nematodes were frozen in liquid nitrogen and RNA extraction was performed as described in the next section.

### RNA extraction and RT-qPCR

PWN RNA extraction was performed for each knockdown treatment (*BXY_079*, *BXY_022* and *gfp*) using the RNeasy Plant Mini kit (QIAGEN) and treated with RNase-free DNase (QIAGEN), according to the manufacturer’s instructions. The quantity and quality of the extracted RNA were assessed by an ND-2000/2000c NanoDrop spectrophotometer (Thermo Scientific, USA) and gel electrophoresis. The cDNA was synthesised using the SuperScript® III first-strand synthesis system (Invitrogen by Life Technologies) following the manufacturer’s instructions. Afterwards, the cDNA was used for RT-qPCR (reverse transcription-quantitative PCR) with LUNA Universal qPCR Master Mix (NEB) to evaluate the effect of RNAi assay. The relative gene expression was normalised using the 2^-ΔΔCt^ method (Livak & Schmittgen, 2001) against two reference genes, actin (Vicente *et al*., 2015) and elongation factor 1-α (Chen *et al*., 2021). The primers used for this analysis are shown in Supplementary Table S1.

### RNA sequencing

The RNA samples, each with an RNA integrity number over the value of 9 (Agilent Bioanalyzer), were sent to sequencing. Nematode mRNA was purified from total RNA and quantified libraries were sequenced on Illumina PE150 platform in Novogene UK, Ltd. After sequencing, adapter trimming and quality filtering were performed using Trimmomatic (v0.39; Bolger *et al*., 2014). Reads with low-quality bases from beginning and ends or with length shorter than 60 bp were discarded. In addition to data generated from this study, publicly available RNA-seq data for *B. xylophilus* life-stages were obtained from the DNA Data Bank of Japan (DDBJ) Sequence Read Archive under study accession DRP002610 (Tanaka *et al*., 2019). The dataset comprised 30 samples with accession numbers DRR029450 through DRR029457 and DRR141203 through DRR141222, representing multiple developmental stages and experimental conditions. Raw sequencing data were downloaded using the SRA Toolkit (v3.0.10).

### Reference Genome and Functional Annotation

The reference genome for *B. xylophilus* (PRJEA64437, WBPS19 release) was downloaded from WormBase Parasite (https://ftp.ebi.ac.uk/pub/databases/wormbase/parasite/releases/WBPS19/species/bursaphelenchus_xylophilus/PRJEA64437/). Protein functional annotation was performed using multiple complementary approaches. Gene Ontology terms and pathway annotations were assigned using InterProScan6 (Jones *et al*., 2014). Custom Python scripts were developed to parse and reformat the GFF3 output into tab-separated files linking protein identifiers to their corresponding GO ontology terms. Signal peptides were predicted using SignalP 6.0 (Teufel *et al*., 2022) with the eukaryotic organism model in slow mode for maximum accuracy. Transmembrane helices were predicted using two complementary methods: DeepTMHMM (v1.0.24; Hallgren *et al*., 2022) via BioLib local execution and TMHMM 2.0c for comparative validation.

### Read alignment and expression quantification

Quality-filtered reads were aligned to the *B. xylophilus* reference genome using STAR (v2.7.11b; Dobin *et al*., 2013). The maximum number of multiple mappings allowed was set to 1, and the maximum number of alignments output per read was similarly restricted to 1, by setting ‘--outFilterMultimapNmax 1’ and ‘--outSAMmultNmax 1’. Quality control of alignment was performed using the mapinsights bamqc module (v1.0; Das *et al*., 2023) and QualiMap (v2.3; Okonechnikov *et al*., 2016). Gene-level read counts were generated using HTSeq-count (Anders *et al*., 2015). FPKM (Fragments Per Kilobase of transcript per Million mapped reads) and TPM (Transcripts Per Million) values were calculated using StringTie (v3.0.0; Pertea *et al*., 2015).

### Expression clustering analysis

Temporal and condition-specific expression patterns were analysed using the ClusterGVis R package (Zhang, 2022) with the soft clustering algorithm. Soft clustering using the Mfuzz algorithm was performed to identify genes with similar expression trajectories across conditions, with the optimal cluster number determined using the getClusters function prior to final clustering. Visualization was performed using line plots and combined line-heatmap representations. Cluster membership and expression profiles were exported as CSV files for downstream analysis.

### Differential expression and visualization

Differential gene expression between knockdown and GFP control conditions was assessed using DESeq2 v1.48 (Love *et al*., 2014). Genes with fewer than 5 counts in at least 3 samples were excluded. The GFP control condition was set as the reference level. Wald tests were used for significance testing. Genes with Benjamini–Hochberg adjusted *P* < 0.05 were considered differentially expressed. For downstream visualization, a minimum absolute log2 fold-change threshold of 0.5 was additionally applied. Volcano plots were generated using EnhancedVolcano (Blighe *et al*., 2018), displaying log2 fold-change against adjusted *P*-values with cutoffs of 0.5 and 0.05, respectively.

### Gene ontology enrichment analysis

Genes were assigned to clusters based on membership values greater than 0.6 from the Mfuzz clustering results, and gene lists for each of the 8 clusters were extracted for enrichment analysis. GO enrichment analysis was performed using TBtools software (Chen *et al*., 2020) with the background gene set comprising all annotated genes in the *B. xylophilus* genome. Enrichment significance was calculated using hypergeometric tests with Benjamini-Hochberg correction for multiple testing using an adjusted p-value threshold of 0.05. Custom R scripts utilizing ggplot2, dplyr, and patchwork packages were developed for enrichment result visualization. All custom scripts and more details are available in https://github.com/chongjing/ProjectPotpourri/tree/main/01.Bursaphelenchus_RNAseq.

### *In situ* hybridisation

Based on the protocol described by Espada *et al*., 2016, *in situ* hybridisation was performed using digoxigenin-labelled probes to determine the spatial expression patterns of candidate genes. Total RNA was extracted from isolates of PWN using the RNeasy Mini kit (QIAGEN) and treated with RNase-free DNase (QIAGEN) before reverse transcription, according to the manufacturer’s instructions. The quantity and quality of the extracted RNA were assessed by an ND-2000/2000c NanoDrop spectrophotometer (Thermo Scientific), and cDNA was synthesised using the SuperScript® III first-strand synthesis system for RT-PCR (Invitrogen by life technologies) following the manufacturer’s instructions. For each candidate gene, a fragment of approximately 200 base pairs was amplified. The primers used for this analysis are shown in Supplementary Table S1. The RT-qPCR amplification was performed Platinum^®^ Taq DNA Polymerase High Fidelity (Invitrogen by Thermo Fisher Scientific, Waltham, MA, USA). PCR products were purified by MinElute® Gel Extraction Kit (QIAGEN) and MinElute® PCR Extraction Kit (QIAGEN). The resulting PCR products were used as a template for generation of sense and antisense DIG-labeled probes using a DIG DNA Labeling Mix (Roche). The nematodes were fixed overnight in 2% (wt/vol) paraformaldehyde and cut using a vibrating aquarium air pump with a razor blade vertically on the slide and moved slowly back and forth across the nematode suspension. Nematodes were pre-treated with proteinase K and hybridised probes within the nematode tissues were detected using an anti-Digoxigenin-AP FAB fragments (Roche) conjugated to alkaline phosphatase detection buffer and its substrate, NBT/BCIP Stock Solution (Roche). Nematode segments were observed using a light microscope (Olympus BX50) and pictures were taken using Cell^D^ 3.2 (Olympus Soft Imaging Solutions). The sense probe was used as a control.

## Acknowledgements

This work is partly funded by the Portuguese Foundation for Science and Technology (FCT) under the Project NemaWAARS - 10.54499/PTDC/ASP-PLA/1108/2021 and individual funding for M. Mendonça PhD fellowship (2024.00901.BD) and M. Espada (10.54499/CEECIND/00066/2018/CP1560/CT0003). The authors acknowledge the R&D unit MED - Mediterranean Institute for Agriculture, Environment and Development (https://doi.org/10.54499/UID/05183/2025) and the Associate Laboratory CHANGE - Global Change and Sustainability Institute (https://doi.org/10.54499/LA/P/0121/2020). Work on plant-parasitic nematodes at the University of Cambridge is supported by DEFRA licence 125034/359149/3, and funded by BBSRC grants BB/R011311/1, BB/S006397/1, BB/X006352/1, and BB/Y513246/1, a Leverhulme grant RPG-2023-001, and a UKRI FrontierResearch Grant EP/X024008/1, the Gatsby Charitable foundation, and through the Cambridge-Africa ALBORADA Research Fund.

## Supplementary files

**Table S1.**
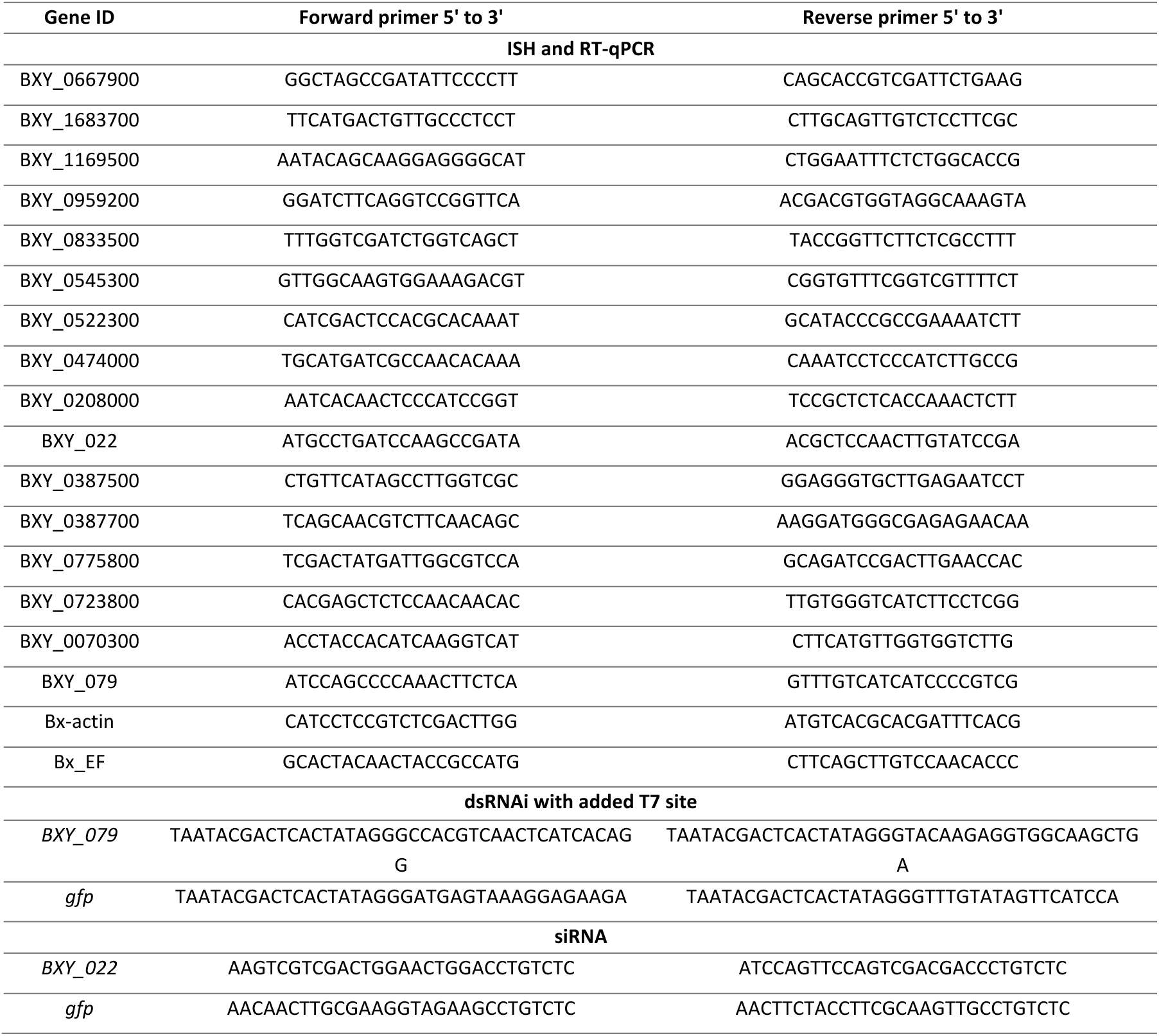
List of primers used in this study.

**Figure S1.**
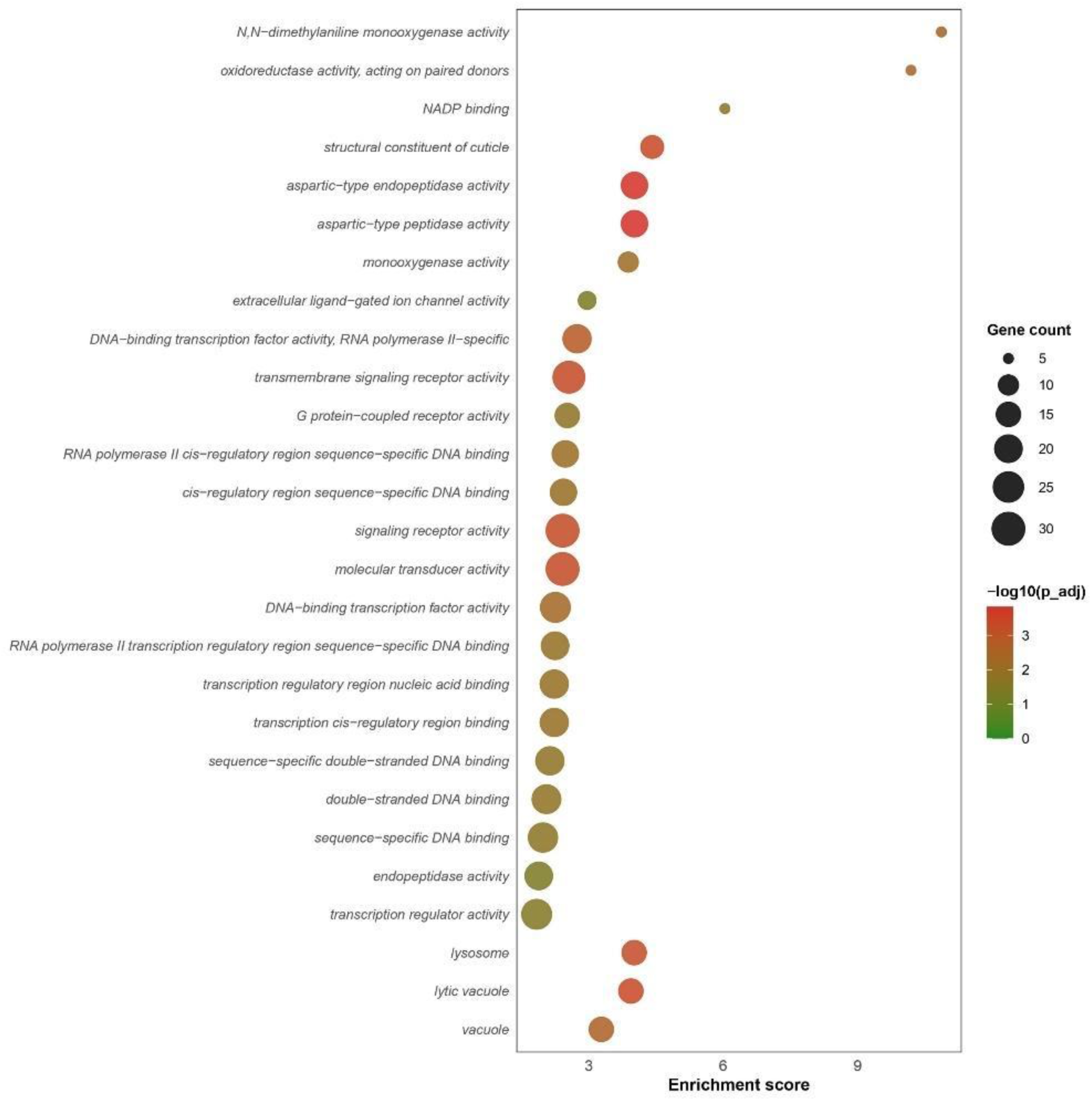
GO term enrichment analysis of *BXY_079*-activated genes.

**Figure S2.**
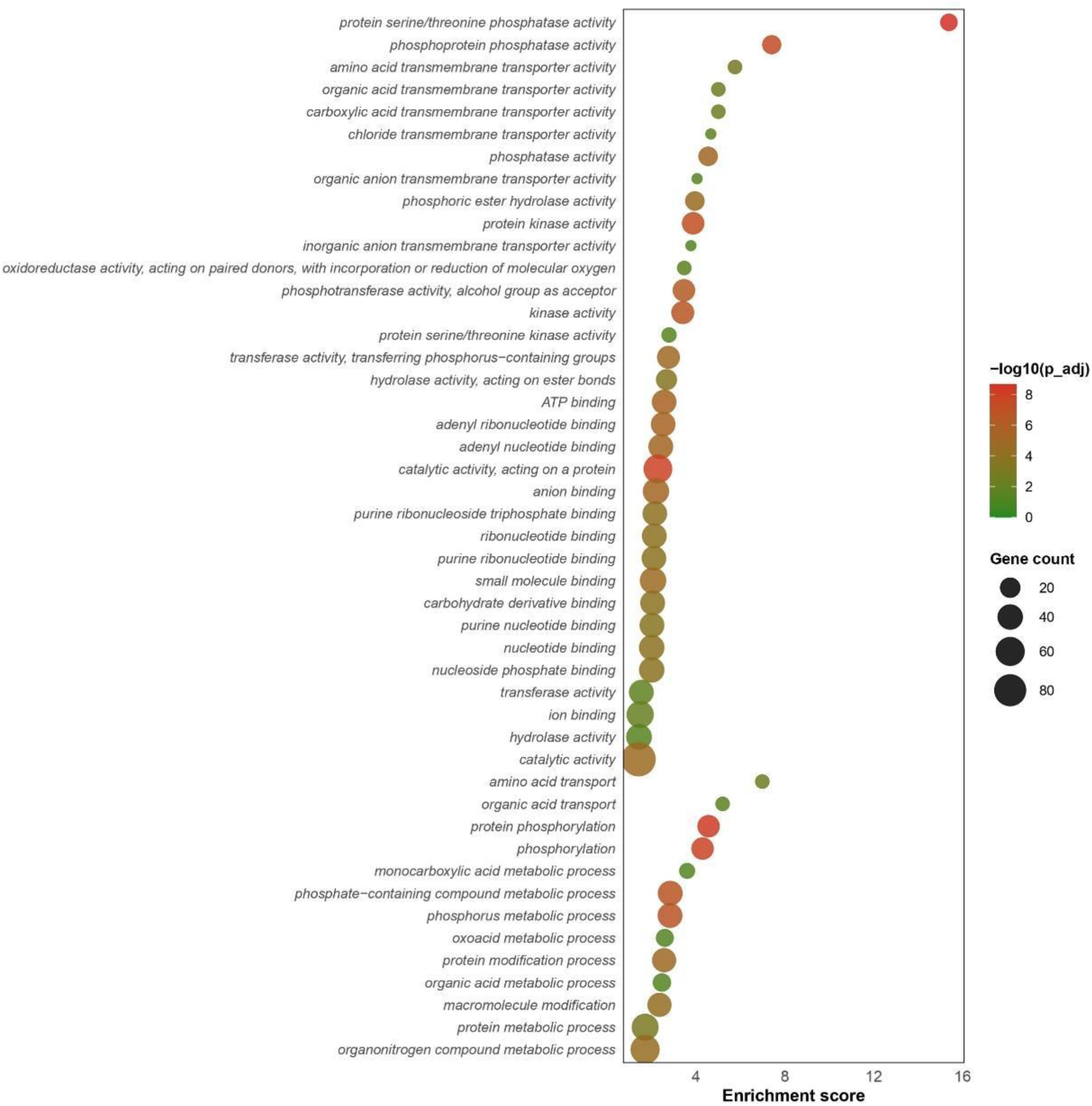
GO term enrichment analysis of *BXY_022*-repressed genes.

